# Genome-wide SNP data reveal recent population structure of *Huidobria fruticosa* (Loasaceae), a paleo-endemic lineage from the Atacama Desert

**DOI:** 10.1101/2023.12.01.569612

**Authors:** Felix F. Merklinger, Yichen Zheng, Tim Böhnert, Federico Luebert, Dörte Harpke, Alexandra Stoll, Thomas Wiehe, Marcus Koch, Dietmar Quandt

## Abstract

The Atacama Desert is a biodiversity hotspot of neo-endemic radiation, where long-term aridity and complex physiographic processes create a unique environmental setting. Current species assemblages are mainly concentrated in highly patchy loma formations, and plant populations occurring in these are often geographically isolated from each other. Despite a general consensus on long-term aridity in the Atacama, climatological and geological evidence points to repeated climate change, making the Atacama Desert an ideal system for studying population genetic processes in highly unstable habitats. We are analyzing the genetic structure within and between populations of *Huidobria fruticosa*, a paleo-endemic lineage of the Atacama Desert, to shed new light on its biogeographic history and broaden our understanding of the evolution of life in extreme aridity, as well as plant evolution in response to a changing environment. To do this, we analyzed SNP data from genotyping-by-sequencing of 354 individuals from 21 populations. Our results suggest that, despite being an ancient lineage, the current population structure of *Huidobria fruticosa* only reflects changing abiotic conditions over the last 2 million years. We therefore conclude that the present distribution, together with the evolutionary processes documented here, is the result of climatic fluctuations and prolonged periods of hyperaridity during the Pleistocene. Building on this understanding, our findings contribute to a global narrative that highlights the complex interplay between climate change and evolutionary dynamics, and emphasize the importance of deserts as living laboratories for deciphering how species have historically adapted to some of the most extreme habitats on Earth.

## INTRODUCTION

On a deep geological timescale, patterns of biodiversity evolve in direct response to geological and climatological processes (e.g., Axelrod, 1967; Gillespie & Roderick, 2014). In extreme environments, such as those marked by prolonged and severe drought, these evolutionary processes are more severely affected than in more stable habitats (such as the wet tropics), because greater spatio-temporal species turnovers may lead to higher extinction rates (Gaston & Spicer, 2004; Stebbins, 1952). Through contraction and expansion of suitable habitats during climatic oscillations, populations may undergo phases of isolation, experience genetic drift, and reconnect to form new species and species assemblages (Stebbins, 1952). Environmental heterogeneity has direct consequences for plant adaptation and as such is reflected in the genetic make-up of the organisms studied.

The Atacama Desert of northern Chile is an extremely arid environment with a hyper-arid core (between 19° and 22°S) receiving less than 1 mm/yr precipitation (Houston, 2006). Nevertheless, precipitation events do occur occasionally, delivering moisture in two major ways: (1) through precipitation events in the Andes leading to occasional discharge into the desert core, and, on a more continuous level, feed the few surface rivers that cross the Atacama on their way towards the Pacific Ocean; (2) regular advection fog that meets the westward facing slopes of the coastal cordillera at a more or less constant altitude due to a temperature inversion (Garreaud et al., 2008). As a result, the Atacama Desert, despite being considered one of the driest places on earth (McKay et al., 2003), harbours a comparatively large number of plant species with a high rate of endemism (Dillon & Hoffmann, 1997). Vegetation is confined to those areas that experience periods of moisture availability, while in contrast, vast stretches of the Atacama Desert are almost totally barren, leaving the populations of many species in geographical isolation over long periods of time (Böhnert et al., 2022; Merklinger et al., 2020, 2021). In addition, the Atacama is an ancient desert with arid conditions persisting at least since the early Miocene (Dunai et al., 2005) and possibly even earlier (Hartley & Chong, 2002; Ritter et al., 2018a). For these reasons, the Atacama Desert is an ideal study area for the evolution of life under hyper-arid conditions.

While a number of botanical and biogeographical studies on the Atacama Desert has recently been published (e.g., Böhnert et al., 2019, 2020; Dillon et al., 2009; Luebert, 2011; Luebert et al., 2009, 2011), very few studies have so far focused on population genetics and plant population structure in relation to geological and climatological events (e.g., Baranzelli et al., 2014; Luebert et al., 2014; Ossa et al., 2013, 2017; Viruel et al., 2012), and even less have studied population dynamics of plants distributed across the hyper-arid core of the Atacama Desert (Merklinger et al., 2020, Koch et al., 2022).

The genus *Huidobria* Gay (Loasaceae, subfamily Loasoideae) is endemic to the Chilean Atacama and currently consists of two taxonomically recognized species, *Huidobria fruticosa* Phil. and *Huidobria chilensis* Gay (Acuña et al., 2017; Grau, 1997; Figure 1). Both species occupy the hyper-arid zones of the Atacama (the definition of hyper-aridity follows Luebert & Pliscoff, 2017). The distribution of the species in focus here, *H. fruticosa*, encompasses the coastal cordillera as well as the western slopes of the Andes. Its populations appear separated from each other by the barren desert pampa that stretches from the eastern slopes of the coastal cordillera eastward to the Andes. Because they are found at either end of the riverbeds (quebradas) that cross the Atacama from the Andes to the Pacific coast, it seems plausible that these (usually dry) rivers act as dispersal corridors for *H. fruticosa*.

**Figure 1:**
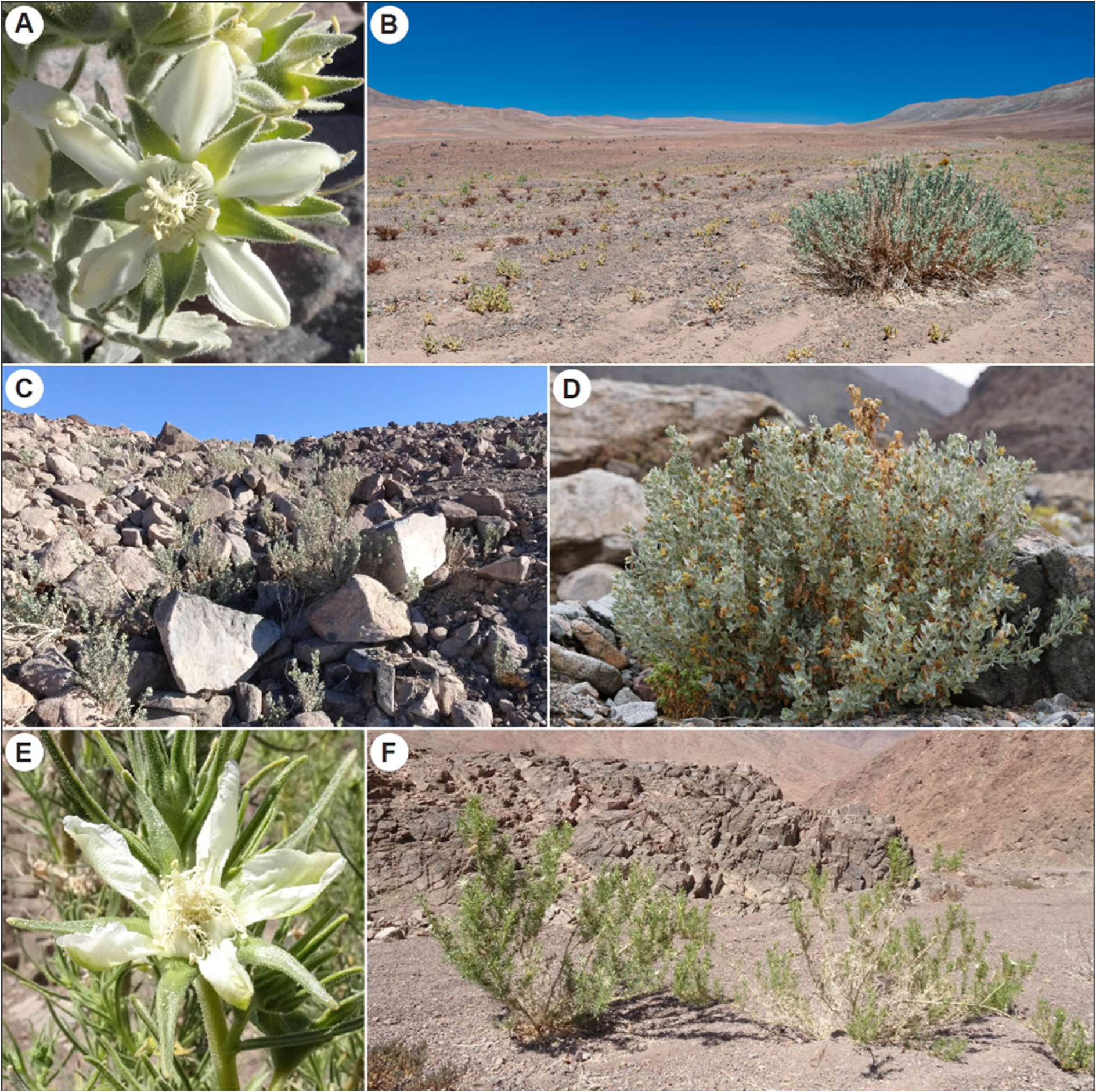
*Huidobria fruticosa* (A-D) and *H. chilensis* (E-F) flowers and habit. **A** Close-up of *H. fruticosa* flower. **B** Typical habitat with old-growth individual in presumably long-term population. **C** Newly appeared population after localized precipitation event. **D** Regrowth from the base of the plant. **E** Flower close-up of *H. chilensis*. **F** Habit of *H. chilensis*. Images: FFM (A, C – F), TB (B).

An ecological peculiarity of this species is its ability to occupy extremely dry habitats where often only few other plants occur. Based on our own field observations, the plants can die back periodically above ground, but re-grow to flowering size in the course of a few months or even weeks (Figure 1D). It is also thought that *H. fruticosa* has the ability to develop a deep root network that eventually reaches into the groundwater layers and is thus able to establish populations that are to a certain degree independent from above ground precipitation availability. Depending on the availability of water, whether above or below ground, plants may persist for longer periods of time, whereas elsewhere they are more ephemeral. We also observed areas covered by many hundreds of individuals where there was barren land a few months before. Such mass occurrence may originate from a seed bank, which in turn can facilitate populations to re-connect from isolation after precipitation events, allowing subsequent gene-flow.

Based on morphological traits, it has been suggested that the evolutionary split of *Huidobria* from its congener as well as closest relatives in the core Loasoideae occurred early in their evolutionary history (Grau, 1997). This has recently been confirmed by molecular clock dating studies, suggesting a divergence between *H. fruticosa* and the only other species in the genus, *H. chilensis*, during the early Eocene, around 40–60 Ma (Acuña et al., 2019). Both lineages could thus be classified as paleo-endemics (Stebbins & Major, 1965), which contrasts several other Atacama groups studied for this region, the neo-endemics – groups that originated more recently in the Atacama Desert, such as *Nolana* (Solanaceae, Dillon et al., 2009), *Heliotropium* (Heliotropiaceae, Luebert et al., 2011), *Malesherbia* (Malesherbiaceae, Gengler–Nowak, 2002) or *Cristaria* (Malvaceae, Böhnert et al., 2019, 2022). Interestingly, the presence of ancient endemic lineages is rare in the Atacama Desert, where predominantly neo-endemic lineages have been found (Scherson et al., 2017), a feature also documented in other deserts (Thornhill et al., 2016, 2017).

Considering *H. fruticosa* as an ancient endemic lineage of the Atacama Desert, we aimed to understand the population dynamics of this species in the light of past climatological processes. We ask if populations have undergone long-term isolation in parallel to the long-lasting aridity of the Atacama Desert. In such a case, we would expect a comparatively high genetic differentiation among populations with respect to other, neo-endemic plant groups. Alternatively, regular precipitation events, accompanied by seed bank effects and/or dispersal events may lead to very little genetic differentiation among subpopulations. To obtain a clearer picture about demographic and genetic substructure, we employed genotyping-by-sequencing (GBS) with 354 samples from 21 populations of *H. fruticosa* and analyzed single-nucleotide-polymorphism (SNP) data. We hypothesize, that the population genetic processes of *H. fruticosa* have been shaped by the interplay of long-term aridity in the Atacama Desert and the divergence time retrieved for this organism.

## MATERIALS AND METHODS

### Study system

*Huidobria fruticosa* is documented in northern Chile from 25.4° S to about 18.2° S (Grau, 1997). Locality information was taken from Grau (1997) and ground-truthed during four field campaigns between 2017 and 2019. In the present study, 21 populations were sampled between quebrada San Ramón near Taltal (25.4°S) and quebrada Tiliviche near the town of Pisagua (19.5°S) (Figure 2). To the north of Pisagua, we could not find the species despite previous reports of it (Grau, 1997). Populations are usually confined to the extremities in the east (Andes) and west (coastal cordillera) of the dry riverbeds that cross the Atacama, with populations at either end of these riverbeds separated by the barren desert pampa. The plants are shrubs of usually about one to two, rarely three meters in height. They produce white flowers that contain small amounts of highly concentrated nectar, indicating insect pollination (Ackermann & Weigend, 2006). Masses of tiny seeds are released from capsules, and as typical dust-flyers are distributed by wind (Weigend et al., 2004).

**Figure 2:**
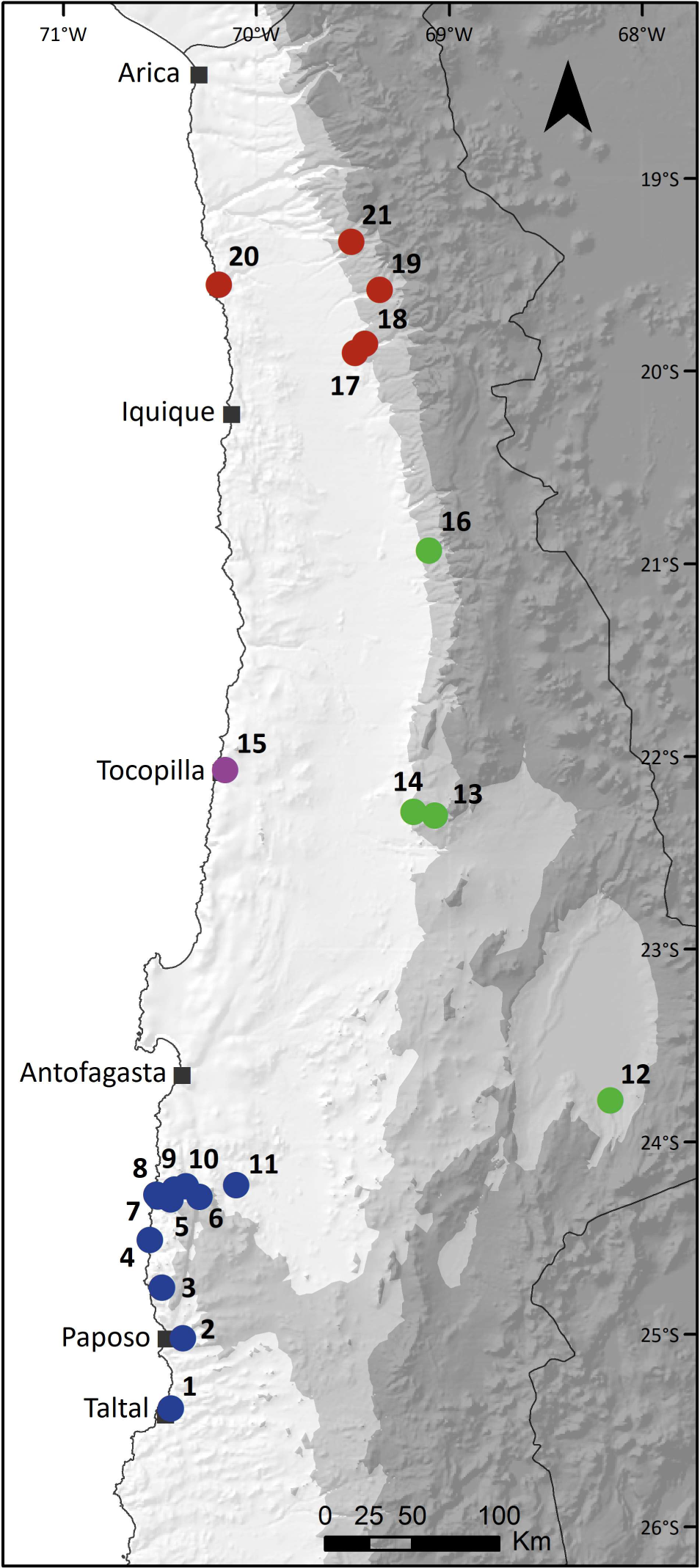
Distribution map of *Huidobria fruticosa* in the Atacama Desert in northern Chile. Each dot represents a sampled population, with the colors corresponding to the geographic population clusters used in the following figures. Map inlet shows South America with a rectangular representing the study area in northern Chile.

### Population sampling

From each of the 21 sampled populations leaf tissue was collected in average from 15 individuals (at least 5 individuals and up to 25 individuals, depending on the population size) and stored in silica gel. Details about each population are given in Table 1. Sampling was carried out along a transect crossing the entire population in order to obtain the best possible coverage. At least one voucher specimen from each population was deposited at the herbarium of the Nees Institute for Biodiversity of Plants, University of Bonn, Germany (BONN) and the herbarium of the University of La Serena, Chile (ULS). A detailed documentation for each population is given in Suppl. Table S1.

**Table 1:** List of *H. fruticosa* populations sampled between 2016 and 2019. Consecutive numbers are given for cross-referencing with the figures, as well as the population field name, location with coordinates in decimal degree (latitude & longitude), individuals sampled per population and genetic cluster based on population analyses.

| Nr | Name | Locality | Latitude | Longitude | Indiv. | cluster |
| --- | --- | --- | --- | --- | --- | --- |
| 1 | HF_2019 | Quebrada San Ramón | -25.38642 | -70.44410 | 14 | South Coast |
| 2 | HF_430 | Lala Kama (Paposo) | -25.01979 | -70.38057 | 20 | South Coast |
| 3 | HF_118 | Cerro Paranal | -24.75786 | -70.48971 | 10 | South Coast |
| 4 | HF_451 | Quebrada Caleta Botija | -24.51074 | -70.55319 | 20 | South Coast |
| 5 | HF_453 | El Cobre to Antofagasta | -24.29918 | -70.44527 | 20 | South Coast |
| 6 | HF_423 | Panamericana km 95 | -24.28712 | -70.29526 | 15 | South Coast |
| 7 | HF_530 | Quebrada El Cobre | -24.28437 | -70.51086 | 5 | South Coast |
| 8 | HF_452 | Above El Cobre | -24.27407 | -70.51920 | 6 | South Coast |
| 9 | HF_532 | El Cobre Mirador | -24.24908 | -70.42477 | 20 | South Coast |
| 10 | HF_533 | El Cobre to Antofagasta | -24.23187 | -70.36576 | 20 | South Coast |
| 11 | HF_615 | SE Antofagasta | -24.22561 | -70.10584 | 15 | South Coast |
| 12 | HF_303 | Tilomonte to Tilopozo | -23.78628 | -68.16624 | 23 | Andean |
| 13 | HF_184 | NW of Chuquicamata | -22.30851 | -69.07521 | 23 | Andean |
| 14 | HF_185 | W of Chuquicamata | -22.28956 | -69.18568 | 20 | Andean |
| 15 | HF_537 | Quebrada Tocopilla | -22.07125 | -70.16314 | 20 | Tocopilla |
| 16 | HF_540 | Quebrada Huatacondo | -20.93127 | -69.10583 | 8 | Andean |
| 17 | HF_110 | Tarapacá | -19.90727 | -69.48951 | 10 | North |
| 18 | HF_661 | Pachica | -19.85838 | -69.43726 | 20 | North |
| 19 | HF_663 | Quebrada Aroma | -19.57878 | -69.36117 | 25 | North |
| 20 | HF_662 | Pisagua Viejo | -19.55212 | -70.19423 | 20 | North |
| 21 | HF_678 | Quebrada Camina | -19.32942 | -69.50792 | 20 | North |

### DNA extraction, library preparation and genotyping-by-sequencing

DNA extraction followed Merklinger et al. (2020). In brief, DNA was isolated from silica-dried leaf material, following the Macherey Nagel Nucleo Mag 96 protocol (Macherey Nagel, Düren, Germany) and processed by a Thermo Fisher Scientific KingFisher Flex benchtop system (Thermo Fisher Scientific, Waltham, MA, United States), binding DNA to NucleoMag C-beads. Quantity and quality of extracted DNA was tested by gel electrophoresis using Lonza GelStar Nucleic Acid Gel Stain (100x) including a sample of 20 ng linear, double-stranded Lambda DNA (New England Biolabs, N3011S). Qubit 2.0 Fluorometer (Life Technologies, Carlsbad, CA, United States) measurements of selected individuals were taken and samples were standardized to 20 ng/μl. For library preparation 200 ng of genomic DNA were digested with the restriction enzymes *Pst*I-HF (New England Biolabs, R3140S) and *Msp*I (New England Biolabs, R0106S). Library preparation, individual barcoding, and single-end sequencing on the Illumina HiSeq 2500 followed Wendler et al. (2014). Barcoded reads were de-multiplexed using the CASAVA pipeline 1.8 (Illumina, Inc.). The obtained raw sequence reads (0.5–4 million per individual) were adapter trimmed and quality trimmed (phred score >25) with CUTADAPT v1.16 (Martin, 2011), and reads shorter than 65 bp after adapter removal were discarded. Sequence reads for the GBS Illumina runs were deposited in the European Nucleotide Archive under the study accession PRJEB62761.

### Assembly parameters

A de novo assembly of the GBS data was carried out using ipyRAD v0.9.71 (Eaton & Overcast, 2020). This assembly contained 354 individuals of *H. fruticosa* and three individuals of *H. chilensis* as an outgroup for the triple comparison method (see section Genetic diversity, past population sizes and sequence divergence), giving a total of 357 individuals. Two different sets of output files were generated through the ipyRAD branching option, one with all 357 individuals, another without the outgroup samples, to be used for population structure analyses. For both branches, the minimal sample number per locus was set to 150, the clustering threshold within samples was set to 0.9. *H. fruticosa* is assumed to be diploid (2n = 36; Grau, 1997), so the ploidy level was set to diploid accordingly. For the other parameters the default settings of parameter files generated by ipyRAD were used. For all downstream analyses, two additional steps of filtering were applied: first all sites (columns) that contained a gap in any individual were removed, then all individuals (rows) that had 6000 or more unresolved nucleotides (N) were excluded. The resulting matrix contained 315 individuals and 21,086 sites.

### Genetic structure

We conducted a principal component analysis (PCA) using 4,118 SNP’s that did not contain any “N” and had only two possible nucleotide states. Homozygotes of the two states were coded “0” and “1” and heterozygotes were coded “0.5” for the input matrix. The first three principal components were plotted and interpreted in the light of population grouping.

Population structure was further explored using the STRUCTURE v2.3.4 (Pritchard et al., 2000) as implemented in ipyRAD v0.9.71, to cluster all sampled individuals into K-distinct populations. Values for K were tested for K = 2–6 with 20 replicates run per test. Each replicate was run for 800 000 Markov Chain Monte Carlo (MCMC) steps with a burn-in period of 200 000.

### F_ST_ and isolation by distance

Average *F_ST_* across all loci was calculated based on pairwise comparisons between populations and visualized in a heat map. Further, we tested for isolation by distance by correlating the average *F_ST_* to geographical distance between each pair of populations using a Mantel test with the R package vegan v.2.5-4 (Oksanen et al., 2019).

### Genetic diversity, past population sizes and sequence divergence

The population genetic parameter θ = 4 Nμ can be estimated with three methods: θw (number of SNPs divided by harmonic number of 2n-1), θπ (average different SNPs between two haploid chromosomes) and θs (number of singletons). Under panmixis (no migration or population structure), without selection and with and constant population size, the three estimates should be equal. θw > θπ > θs can result from recent population size reduction or internal structure, while the opposite can be caused by recent population size increase or external immigration. Based on the SNP matrix, we calculated three estimates of the population-scaled mutation rate θ, a measure of genetic diversity that is independent of sample size. θπ is the mean number of differences between two haploid genomes in the population. θw is the total number of variable sites in the population divided by the harmonic number of 2n-1 (Watterson’s estimator). Finally, θs is the number (raw count) of singletons, polymorphic sites, where one allele is present only in one genomic copy in the entire genome. These values are aggregated across the entire genome and presented as the total amount of diversity in the SNP matrix.

To be able to compare genetic diversity (π) among populations, and also with other species, we determined π also on a per nucleotide scale. We used the program Stacks v2 (Catchen et al. 2013) with standard parameters for de novo analysis and with the wrapper script "denovo_map.pl" (see https://catchenlab.life.illinois.edu/stacks/manual/). As input we used only those quality filtered FastQ reads, which could be trimmed to a size of exactly 100 bp. Data on genetic diversity are given in Table 2.

**Table 2:** List of population statistics for each population based on STACKS output. Consecutive numbers are given for cross-referencing with the figures and Table 1.

| Nr | Sites | Variant Sites | Polymorphic Sites | % Polymorphic Loci | Pi |
| --- | --- | --- | --- | --- | --- |
| 1 | 1123990 | 14217 | 2977 | 0.2649 | 0.0009 |
| 2 | 1004571 | 12902 | 4648 | 0.4627 | 0.0013 |
| 3 | 1032171 | 12362 | 3390 | 0.3284 | 0.0011 |
| 4 | 1275862 | 16129 | 6887 | 0.5398 | 0.0015 |
| 5 | 1320486 | 16497 | 7659 | 0.5800 | 0.0016 |
| 6 | 1305062 | 16135 | 5953 | 0.4562 | 0.0014 |
| 7 | 1047791 | 13356 | 3737 | 0.3567 | 0.0014 |
| 8 | 1232682 | 15379 | 4485 | 0.3638 | 0.0014 |
| 9 | 1219180 | 15378 | 6670 | 0.5471 | 0.0015 |
| 10 | 1418012 | 17337 | 7890 | 0.5564 | 0.0015 |
| 11 | 1245816 | 15654 | 5633 | 0.4522 | 0.0014 |
| 12 | 1145564 | 14570 | 2807 | 0.2450 | 0.0007 |
| 13 | 889895 | 11456 | 3051 | 0.3429 | 0.0009 |
| 14 | 1215831 | 15405 | 4351 | 0.3579 | 0.0009 |
| 15 | 1326879 | 15984 | 3593 | 0.2708 | 0.0007 |
| 16 | 1361823 | 16472 | 1498 | 0.1100 | 0.0004 |
| 17 | 1259038 | 14979 | 2035 | 0.1616 | 0.0005 |
| 18 | 1290490 | 15492 | 2383 | 0.1847 | 0.0004 |
| 19 | 1069226 | 13260 | 2030 | 0.1899 | 0.0004 |
| 20 | 1410921 | 16182 | 2395 | 0.1698 | 0.0004 |
| 21 | 1213817 | 14687 | 2116 | 0.1743 | 0.0004 |

The program Stairway plot v2.1 (Liu, 2020; Liu & Fu, 2015) was used to estimate the past population sizes from the allele frequency spectra. Because no known estimates of generation time and unscaled mutation rates are available, the results are displayed as if the mutation rate is 1.2e-8 mutations per site per generation, and generation time is one year. The mutation rate is a default value provided by the program. The generation time is arbitrarily decided (default value of 24 years). Only sites that are not “N”s in any individual in that population are counted for both the total site number and the site frequency spectrum.

In order to estimate the time of sequence divergence between population clusters, we developed a “triple comparison method”. For this, the loci were first aligned to the outgroup (OG), *H. chilensis*. To estimate the divergence time between two clusters, we took one individual from each cluster (H1 and H2) and counted the number of sites that were non-ambiguous in both individuals and the OG. Then we counted the number of transitions and transversions between H1 and H2, H1 and OG, as well as H2 and OG. Heterozygotes were counted as half a mutation. For example, A/A and A/G is separated by 0.5 transitions, while A/A and C/G is separated by 0.5 transitions and 0.5 transversions. Following this, we calculated the Kimura-2-Parameter distance (d) between each pair, and the divergence time between H1 and H2 was estimated as t_H1-H2_ = t_0_ × 2 × d_H1-H2_ / (d_H1-OG_+d_H2-OG_), where t_0_ is the divergence time between *H. fruticosa* and *H. chilensis* (∼ 50 Ma, Acuña et al., 2019). Four individuals from four different groups of populations were chosen for each cluster based on the least number of “N”s, except for population 15 (Tocopilla), where the four individuals were from the same (only) population. Therefore, for each pair of clusters the triple comparison was repeated 16 times and the mean estimate was used.

## RESULTS

### Assembly statistics

The assembled data set including the *H. chilensis* outgroup and with 29.07% missing data consisted of 13,400 loci with 40,753 SNPs, of which 31,045 were parsimony informative sites. The second data set (“SNP matrix”) excluding the outgroup samples had 31.66 % missing data and contained 13,405 loci with 35,253 SNPs of which 26,252 were parsimony informative sites. After filtering, 315 individuals and 21,086 SNP sites remained for further analyses.

### Genetic structure

PCA axes 1 & 2 explained 45.8 and 11. 9 % of the total variability, respectively (Figure 3). A first split separates a northern cluster from the rest along the first PCA axis. Along the second axis, a further split between the southern coastal populations and the Tocopilla + Andean populations was observed, while the southern coastal populations appeared closer to the northern populations. PCA axes 1 & 3, explaining 45.8–5.6 % of the total variability, respectively, retrieved the Tocopilla population farthest away from the rest. To further clearly distinguish the four distinct geographical clusters, they are from here on referred to as NC (northern cluster), AC (Andean cluster), SC (southern cluster) and TC (Tocopilla-cluster).

**Figure 3:**
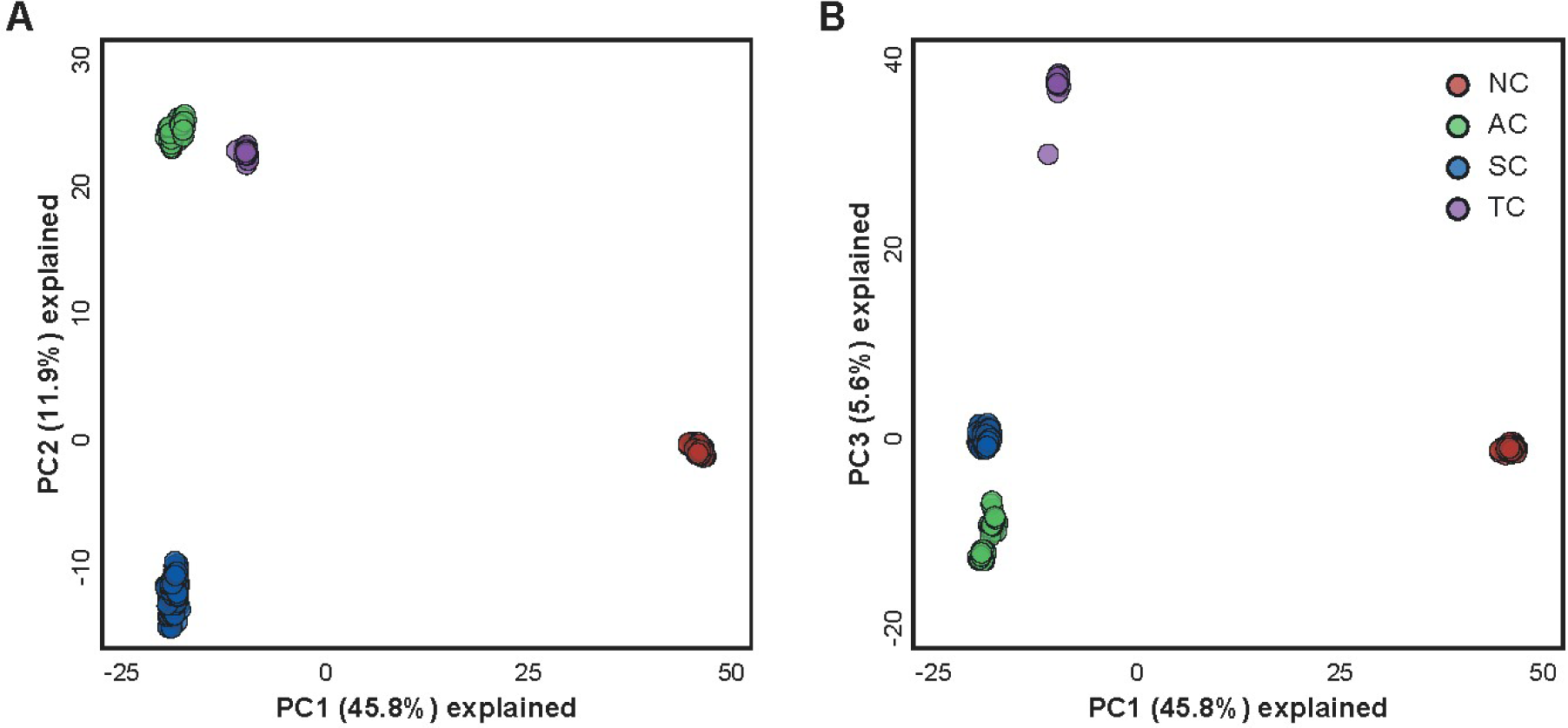
Principal component analysis with randomly sampled, unlinked SNPs. Each dot represents one individual; color-coding represents the four main clusters identified (NC = Northern Cluster, AC = Andean Cluster, SC = Southern Cluster, TC = Tocopilla Cluster).

Results of the structure analysis (Figure 4) are congruent with the results from the PCA, but without a clear separation of the TC population. The optimal delta K value was K = 3 followed by K = 5. At K = 2 the NC, including all populations north of Iquique at 20°S (populations 17–21), separated from the rest. At K = 3 the AC separated from the NC and SC, containing populations 12–16. At K = 5, the TC (population 15) separated from the other populations, but retaining a weak connectivity to its closest Andean neighbor, population 16 near the locality of Huatacondo and population 11, the northern most population of the SC.

**Figure 4:**
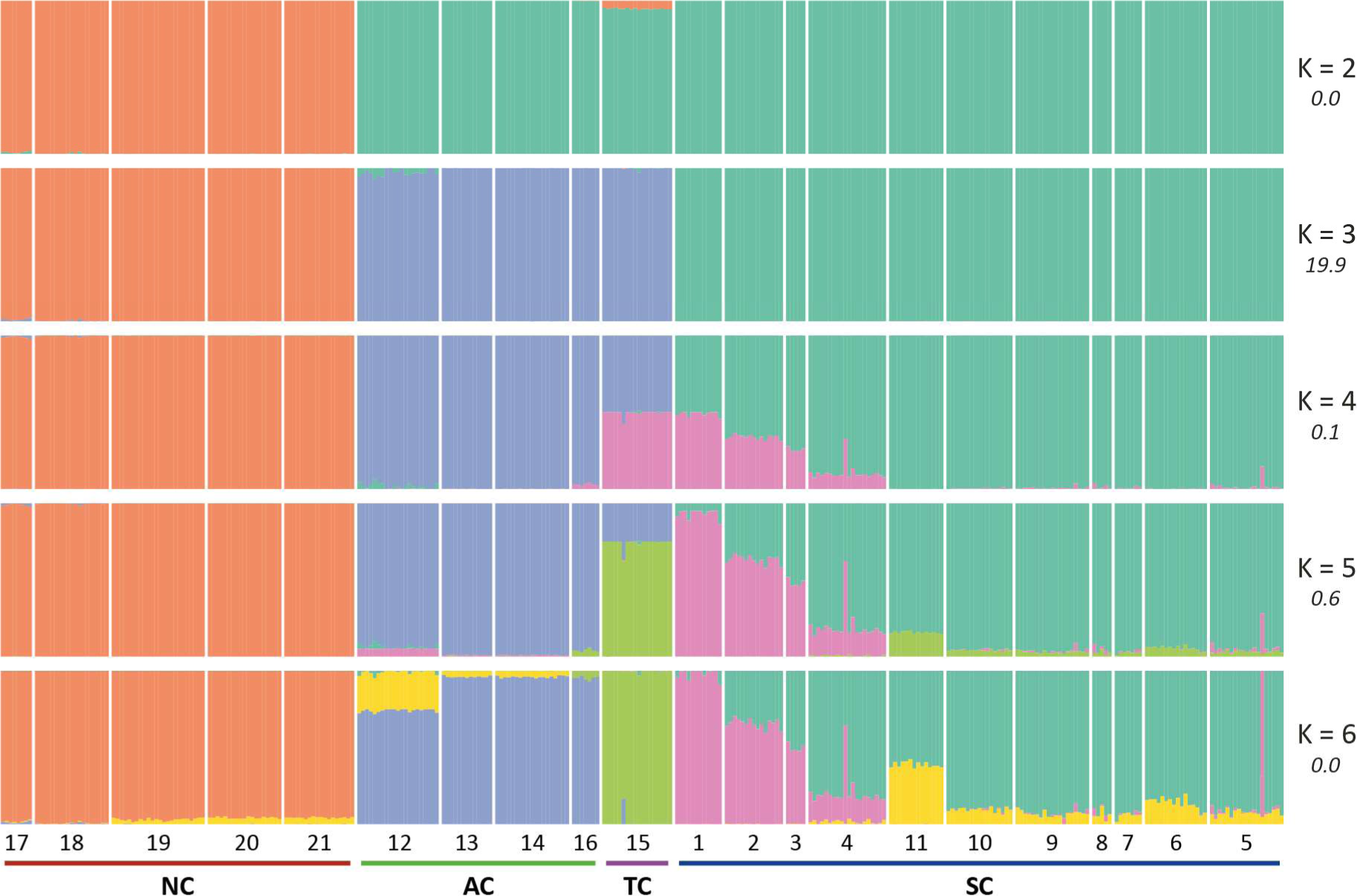
Population structure of the 21 sampled populations of *H. fruticosa* grouped in four geographic clusters. Individuals are grouped by their respective population and population numbers are given below (see figure 2). ΔK and the respective K-values are given on the right. The sampled populations (NC = Northern Cluster, AC = Andean Cluster, SC = Southern Cluster, TC = Tocopilla Cluster) are best grouped into three populations (K = 3), while the Tocopilla (TC) population is recognized as a genetically differentiated population under K = 5.

### F_ST_, Mantel test and isolation by distance

Genetic distances between populations were small to moderate (0.05–0.35), however, some distinct tendencies could be observed between the populations belonging to the four geographical clusters (Figure 5). The populations belonging to the NC all shared a very low *F_ST_* among each other (≤ 0.1), as did the populations of the SC. The two Andean populations (13 and 14) equally shared a very low *F_ST_* value (0.05) although this increased with geographical distance toward the other populations belonging to the AC further north (populations 12 and 16; ∼ 0.1). The largest *F_ST_* values were found between the Andean population 16 near Huatacondo and the populations of the NC (0.35). Further, between two of the southernmost populations (1 and 3) and the NC (0.3–0.35), and between the TC (15) and the rest (0.3–0.35). The Mantel-test showed a positive correlation (r = 0.749; P < 0.001) between the geographic distance of the population to each other and the respective *F_ST_* values, indicating that geneflow decreases with increasing distance.

**Figure 5:**
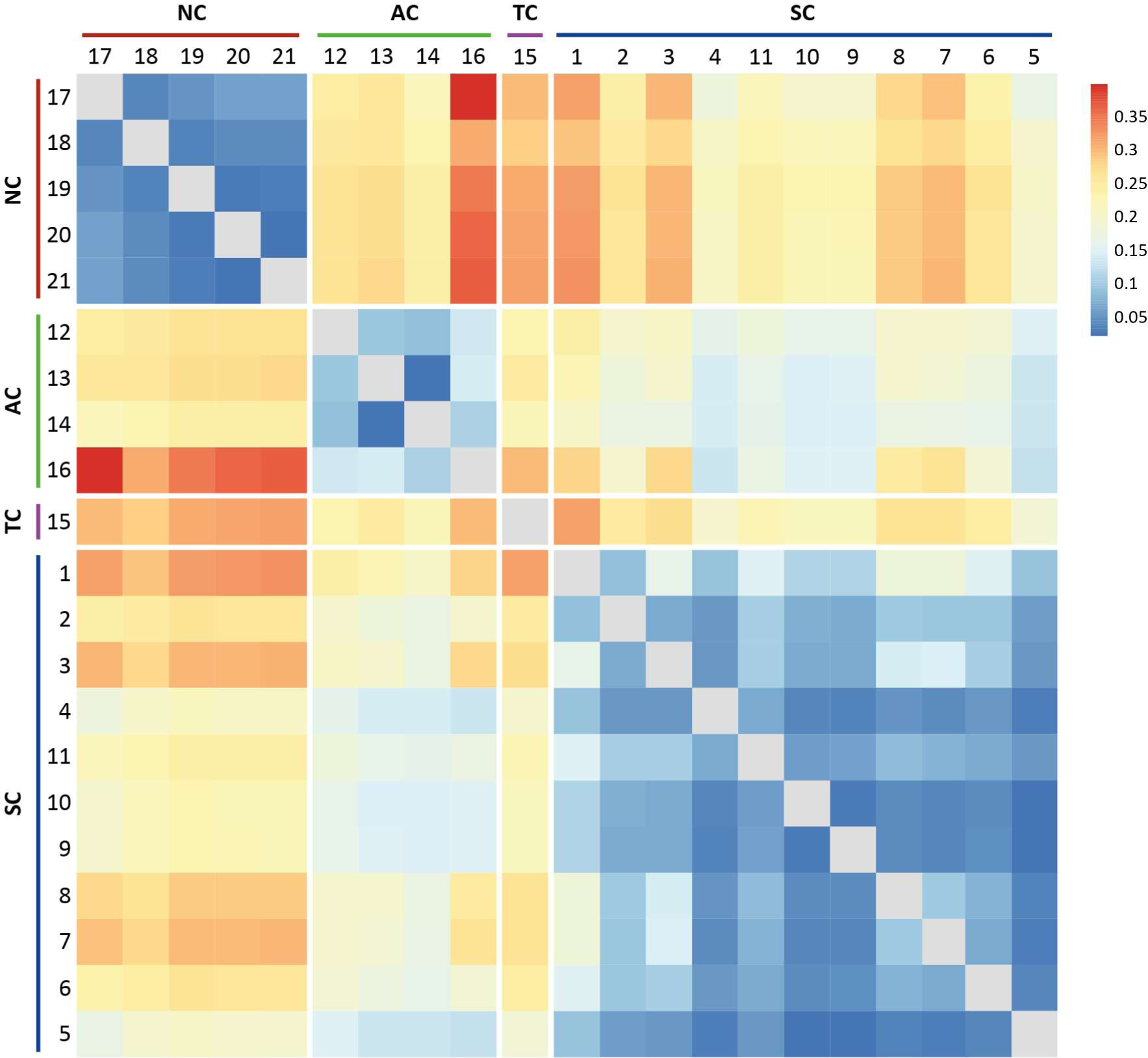
Heatmap of pairwise comparison the average *F_ST_* values across all loci between the 21 populations. High *F_ST_*-values are given in red indicating low gene flow, while low *F_ST_*-values are given in blue indicating high gene flow among populations (NC = Northern Cluster, AC = Andean Cluster, SC = Southern Cluster, TC = Tocopilla Cluster).

### Genetic diversity, past population sizes and sequence divergence

Populations belonging to the NC showed relatively homogeneous estimates of θ, both among subpopulations and among different estimators (θw θπ and θs), supporting the notion of populations being in, or close to, mutation-drift equilibrium (Figure 6). The only exception is population 18, which has an excess of singletons. They are not concentrated in a single individual. Therefore, an initial suspicion that this may be due to a sequence artefact or misclassification was not confirmed. Generally, AC has a larger population size than NC, and appears to be slightly expanding. The only exception here is population 16 (Huatacondo), which has a small Ne, but very even estimates of θ, indicating again that it is close to mutation drift equilibrium, and has stably persisted as a small subpopulation. SC has the largest populations, but most of them, with the exception of populations 4, 5 and 11, appear to be shrinking. Lastly, population 15 (Tocopilla) is intermediate in size and stable. Note the parallel profiles and the close agreement (up to scaling) of the aggregate values of θπ with the per nucleotide estimates of π are given in Figure 6 (first set of histograms and interpolated curve).

**Figure 6:**
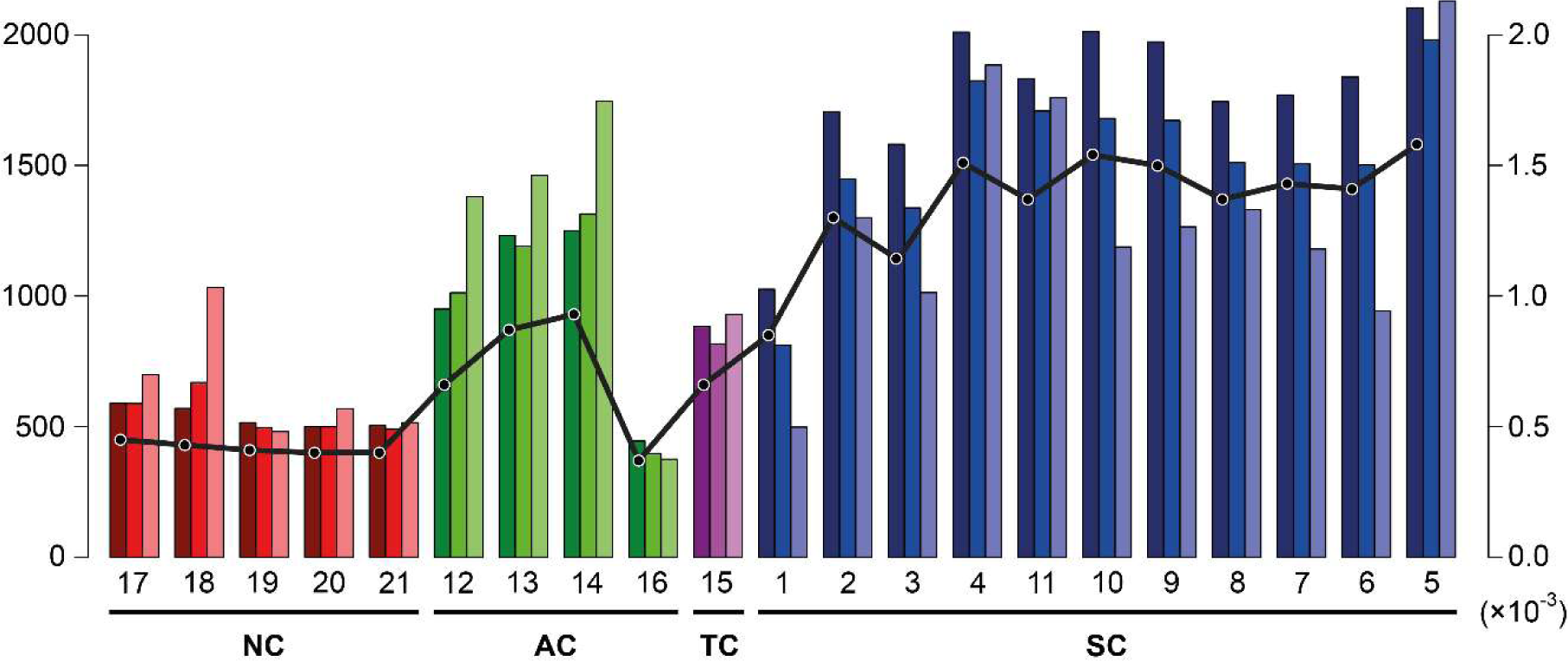
Estimations of aggregate θ (left ordinate) and of genetic diversity π (right ordinate). Dark, medium and light colors refer to the three estimates θπ, θw and θs, respectively. All estimates of θ are based on the total number of polymorphisms seen in the respective populations. Note, that comparability of θ among populations is limited due to varying amounts of missing data in different populations. In contrast, genetic diversity π is drawn on a per nucleotide scale (right y-axis), admitting comparison across populations (and also with other species). Populations are grouped (NC, AC, TC, SC) and labelled as in Fig 2.

The result of the stairway analysis was generally similar to the interpretation of the values of θ: recent reduction in SC, expansion and AC and general stability in NC and TC (Suppl. Figure S1). There was a noticeable decline in all populations belonging to the SC at ∼ 2000 time units, but only a few of these populations showed a subsequent rebound. Also, population 15 (Tocopilla) appeared to have experienced a bottle neck around the same time.

The models from our triple comparison dating output analysis suggest a divergence time of approximately ∼2 Ma for the first split between the NC and the remaining populations. This also corresponds to the split along the first PC axis in the PCA graph (Figure 3). At approx. 1.2–1.4 Mya the AC, SC and TC split further.

## DISCUSSION

### Pleistocene origin of extant Huidobria populations

At the outset of this study, we hypothesized that the population structure of *H. fruticosa* might reveal the signature of long-term aridity of the Atacama Desert. Phylogenetically, the two species of *Huidobria* are separate lineages of Eocene origin (Acuna et al., 2019) and thus pre-date most other Atacama Desert lineages for which dated phylogenies are available (e.g., *Cristaria*, Böhnert et al., 2022; *Malesherbia*, Gengler–Nowak, 2002; *Oxalis*, Heibl & Renner, 2012; *Heliotropium*, Luebert et al., 2012; *Nolana*, Dillon et al., 2009; *Zygophyllaceae*, Böhnert et al., 2020). However, our results suggest that the history of the sampled extant populations is rather recent, dating back only to the last ∼2 Ma. As such, our analyses do not provide insight into the pre-Pleistocene history of *Huidobria fruticosa* in the Atacama Desert. However, we are able to provide a detailed picture of a much more recent population history, providing for the first time the population history of a paleo-endemic plant species from one of the world’s driest deserts.

The here studied populations of *H. fruticosa* are characterized by a well-defined genetic structure which corresponds to four geographically distinguishable population clusters (NC, SC, AC, TC). Our triple comparison analysis suggests an initial split into a northern and southern cluster of populations approximately 2.03 Ma. At about 1.45 Ma, the southern cluster then split further into the Andean (AC) and southern coastal cluster (SC), followed by a third split between the AC and the TC cluster approximately 1.21 Ma. It is to note, however, that this timescale is probably much older than anything that can be retrieved from the within-population variation data, such as the stairway or theta analyses. Because the stairway timescale is always relative to the generation time, the absolute time scale may vary considerably, depending on the actual generation time. Within-population variation data provides information up to 4Ne generations ago; however, in order to obtain Ne we need an estimate of absolute mutation rate.

The Plio–Pleistocene saw a fluctuating climate, with several pluvial phases interspersed with phases of prolonged aridity (Jordan et al., 2014; Ritter et al., 2018b). Cosmogenic nuclide exposure dating to investigate ancient shoreline terraces of the Quillagua-Llamara Soledad Lake demonstrated the presence of several pronounced arid phases 2.65± 0.15 Ma and 1.27 ± 0.47 Mya, when no lake existed (Ritter et al., 2018b), a finding similar to that obtained by Jordan et al. (2014). The dates obtained by Ritter et al. (2018b) fit remarkably well with the dates, which we obtained for our extant population clusters, suggesting a climatically induced shift in the population structure of *Huidobria fruticosa*. This finding is congruent with that made for other plant genera such as *Eulychnia* (Merklinger et al., 2021) or *Cristaria* (Böhnert et al., 2022), with the difference that those groups seem to have diversified in response to climatic shifts, while in *H. fruticosa* this does not appear to have been the case. The lack of observable diversification is surprising considering the phylogenetic age of this lineage. The two known species, *H. fruticosa* and *H. chilensis* are genetically only distantly related (last common ancestor at approx. 50 My; Acuña et al., 2019) and morphologically very different (c.f. Figure 1). While the reasons for such a lack of diversification go beyond this study, similar patterns are also known from other large shrubs or small trees in the Atacama Desert (e.g., Zygophyllaceae: Böhnert et al., 2020). One explanation may be the higher extinction rates in arid landscapes (Stebbins, 1952) and the associated greater species turnover – or simply put: perhaps the past saw a very diverse assemblage of *Huidobria* species similar to other groups today such as e.g. *Nolana* (Solanaceae; Dillon et al., 2009).

During pluvial phases, *H. fruticosa* may have been more widely distributed, possibly forming extensive metapopulations similar to their mass-occurrences today shortly after precipitation events. With the onset of prolonged phases of aridity, however, habitat and populations became increasingly fragmented and populations geographically isolated, in turn leading to the genetic structure of populations that we see today. Similar patterns have been proposed for ancient lineages in other parts of the world, such as *Welwitschia* Hook. from Namibia and Angola, which also shows strong signals of population structure, and which has been interpreted as the result of aridification (Jacobson & Lester, 2003), although in this case, a temporal framework in correlation to climate history is still needed.

### Dispersal and genetic connectivity between Huidobria populations

The three to four geographically and genetically distinguishable *Huidobria* population clusters show varying degrees of gene flow among each other across the hyper-arid landscape. The structure analysis points to continuous gene flow between coastal and Andean populations within the NC, while the SC is genetically isolated from the populations of the Andes, as is, at least to some extent, the TC. This implies that gene flow between the northern coast and the northern Andes has been more continuous than between the Andes and the coast further south. This may in part be explained by the more continuous geography between the coast and the Andes in the far north, facilitating plant dispersal across it. Since the estimated effective population size is very small in this cluster (NC), the excess of singletons could be due to a very recent expansion after a strong bottleneck.

The genetic diversity analysis recovered the largest number of singletons (θs) in the SC, where *F_ST_* values between coastal populations are relatively high. Interestingly, with the exception of populations 9, 11 and 17, the coastal populations show a decreasing trend in the number of singletons obtained from the θ analysis, as well as steep drops in the stairway plot line, possibly indicative of a population bottleneck. A possible explanation may be that, given the highest genetic diversity in the SC, *Huidobria* populations have managed to persist at coastal localities throughout their long-term evolutionary history, and from here, repeatedly colonized different regions of the Atacama Desert.

The coastal terrain of the southern Atacama is strongly fragmented and occupied by a patchy loma vegetation, which is mostly concentrated on the western slopes of the coastal cordillera (Dillon & Hoffmann, 1997; Rundel et al., 1991). The steep coastal cordillera and the hyper-arid pampa del Tamarugal without the Andean-coastal canyon connection may present the species with much stronger climatic and geographical barriers (Ruhm et al., 2020) and so hamper *Huidobria* dispersal, resulting in the Andes-Coast genetic differentiation between populations observed here. The only exception to this is the Tocopilla population, which shows a slight Andean genetic influence. In fact, this population seems more closely related to the Andean population near Huatacondo (population 16) than to any other, geographically closer Andean populations (e.g., populations 13 & 14) in our sampling.

*Huidobria fruticosa* very effectively disperses its seeds, so it appears plausible to infer dispersal events and founder effects at various periods in time as a cause for current population distribution. Effective dispersal and colonization have been shown for other lineages in the Atacama such as, e.g., *Tillandsia* (Merklinger et al., 2020), *Hoffmannseggia* (Simpson et al., 2005) or members of the Boraginaceae (Simpson et al., 2017). This would imply the presence of a source pool, or at least the continuity of populations over time in some areas of the Atacama Desert, from where the repeated colonization of adjacent areas could have taken place. This source pool might be found in the build-up of a soil seed bank by the species’ wind dispersed seeds where they endure arid phases and establish new populations following precipitation events. This would also explain our observation of populations at different localities where previously none were found. Populations would not "move" but establish at fast speed from soil seed banks after precipitation. Effective dispersal further mitigates the extinction risk of newly founded small subpopulations (Den Boer, 1968; Pisa et al., 2019), which is particularly high in arid environments (Stebbins, 1952). Despite effective dispersal, however, *H. fruticosa* has remained an endemic of the extreme north of Chile and has apparently not managed to occupy adjacent regions such as southern Peru, despite similar levels of aridity there. Although there is a floristic break between the coastal floras of southern Peru and northern Chile (Pinto & Luebert, 2009; Ruhm et al., 2022; Rundel et al., 1991), other taxa which have a coastal as well as Andean distribution have migrated across this barrier along the Andean cordillera (Beddows et al., 2017; Dillon et al., 2009; Gengler–Nowak, 2002, Böhnert et al., 2022), which has been proposed to function as a dispersal corridor for taxa (Luebert & Weigend, 2014; Moreno et al., 1994). The absence of *Huidobria* in southern Peru is thus surprising. While this may simply be a sampling bias, a further plausible explanation may be the presence of another Loasoid lineage (*Presliophytum* (Urb. & Gilg) Weigend) in southern Peru with very similar ecological and physiological traits as *Huidobria* in terms of habitat preference (Acuña et al., 2017; Acuña & Weigend, 2017) and seed dispersibility (Weigend et al., 2004). *Presliophytum* may thus act as an effective competitor preventing *Huidobria* to colonize habitats successfully in southern Peru.

### Paleo-endemic desert lineages with recent radiations

Deserts tend to be populated by lineages of recent origin (see e.g., Baldwin, 2014; Byrne et al., 2020) with high levels of neo-endemism (Scherson et al., 2020). Conversely, paleo-endemics are associated with low-latitude and stable climates, where species have survived and relatives gone extinct elsewhere (Daru et al., 2020). Paleo-endemics are thus rare in hot deserts, characterized by climatic instability, with *Welwitschia* being the classic example of an exception to this rule. *Huidobria fruticosa* is another rare example of desert paleo-endemism, though we do know very little about the past history of this lineage. Its genetic population structure, however, does not appear to show any signature of its presumed long evolutionary history. Rather it appears to be the result of recent processes that favored both isolation and connectivity among populations, with gene-flow taking place along the streams between the coastal and the Andean ranges, probably associated with periods of increasing rainfall in recent times. Furthermore, another important prerequisite to explain the here observed population structure is the ability to effectively disperse seeds and to build-up a soil seed bank, and thereby establishing and maintaining isolated populations across the hyper-arid desert landscape. However, north to south gene-flow must be regarded as much more limited, which is possibly a result of large, permanently hyperarid zones between the NC and the other populations that make the survival of any species nearly impossible. In summary, the present-day distribution of *Huidobria fruticosa* can be explained by the impact of abiotic factors, mainly the climatic fluctuation and the prolonged phases of hyper-arid conditions during the Pleistocene. Thereby, abiotic factors account for much biotic diversity in the region because of population isolation and secondary contact leading to diversification of organisms, while biotic diversity also largely depends on the individual life strategy and subsequent carrying capacity of the respective habitats.

## ACKNOWLEDGMENTS

We thank Claudia Schütte (Nees Institute, Bonn), Axel Himmelbach and Susanne König (IPK Gatersleben) for guidance and support with the laboratory processes. We would also like to thank Dr. Julius Jeiter (Nees Institute, Bonn) for feedback and help with preparing figures, as well as Prof. Dr. Maximilian Weigend (Nees Institute, Bonn) for helpful insights into the ecology of the study group as well as substantial feedback on an early draft of the manuscript. This study was funded by the German Research Foundation (DFG) – Project Nr. 268236062 – SFB 1211, Earth: Evolution at the dry limit (http://sfb1211.uni-koeln.de/).

## Data accessibility and Benefit-Sharing

GBS sequence reads are accessible at the European Nucleotide Archive under the study accession PRJEB62761.

The genetic material used in this study is not subject to the regulations of the Nagoya Protocol on Access to Genetic Resources because Chile, the country of origin of these resources, is not a party to the Nagoya Protocol.

## Author contributions

FFM, YZ, TB, TW, DQ & FL designed this study. FFM, TB, FL, AS & DQ collected the samples. FFM & DH generated the sequences. FFM, YZ, TB, TW & FL analyzed the data and interpreted the results. FFM wrote the first draft of the manuscript with subsequent support from YZ, TB, TW & FL. All authors provided critical revision to the drafts of the manuscript.

## Supplementary Material

**Suppl. Table S1**: Population sampling and voucher information for the study organism (*Huidobria fruticosa*) and outgroup (*Huidobria chilensis*).

**Suppl. Figure S1:**
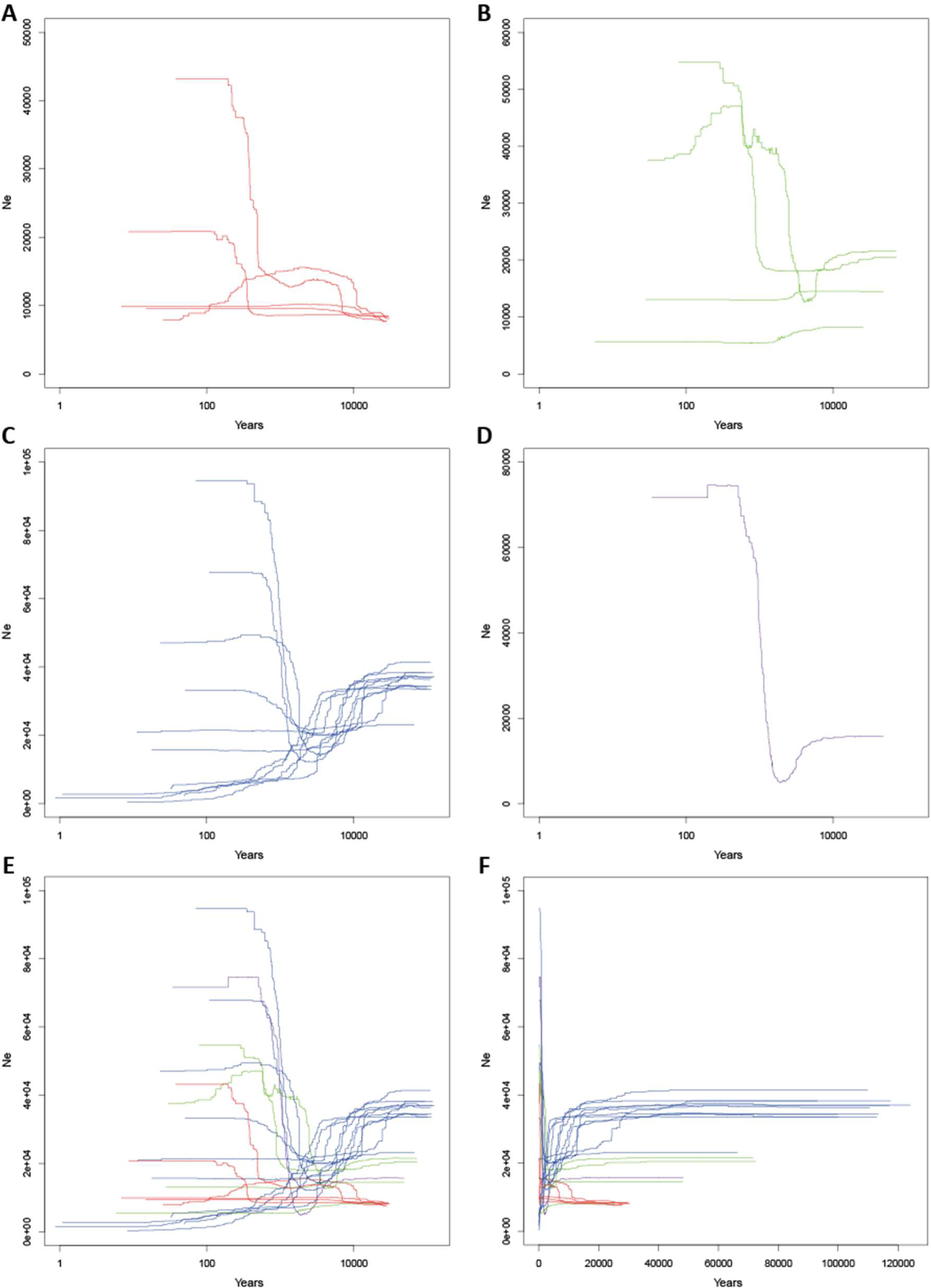
Stairway plots depicting past population sizes calculated from the allele frequency spectra for the individual clusters (A – D) and for all populations together (E – F). **A** Northern cluster. **B** Andean cluster. **C** Southern cluster. **D** Tocopilla cluster. **E** Full plot with all population over 120,000 years. **F** full plot with all populations on a logarithmic scale. Ne stands for effective population size.

